# Regulatory B cells contribute to the alleviation of colitis induced by Dextran Sulphate Sodium after *H.pylori* infection

**DOI:** 10.1101/305094

**Authors:** Xia Li, Jiang Tan, Feng Zhang, Qian Xue, Ning Wang, Xu Cong, Jingtong Wang

## Abstract

**Background:** Epidemiological studies showed that there was an inverse relationship between *Helicobacter pylori* infection and the incidence of inflammatory bowel disease. Our previous research indicated that the regulatory immune responses induced by *H.pylori* infection were not limited to gastric mucosa, and the balance of intestinal mucosal immunity was influenced. In this study, we investigated the role of the regulatory B cells in the effects of the *H.pylori* infection on acute and chronic colitis induced by dextran sulphate sodium.

**Methods:** Mice were infected with *H.pylori* and then colitis was induced by 3% DSS, the CD19^+^IL-10^+^Breg cells and CD4^+^CD25^+^Foxp3^+^Treg cells in blood、 spleen、 mesenteric lymph nodes (MLN)、 Peyer’s Patches (PP) and gastrointestinal mucosa were measured and examined. The anti– and pro-inflammatory cytokines were also detected at mRNA level.

**Results:** In the acute and chronic colitis groups, DAI and colonic histological scores reduced significantly and colon length shorted less; the anti-inflammatory cytokines mRNA expression upregulated while the pro-inflammatory cytokines mRNA expression downregulated in colonic mucosa; and the percentages of CD19^+^IL-10^+^Breg cells in blood spleen MLN、 PP were higher in the *H.pylori*/DSS co-treated group compared with the DSS treated group.

**Conclusion:** *H.pylori* infection can alleviate the acute and chronic colitis induced by DSS. CD19^+^IL-10^+^Breg cells may play a critical role in the alleviation of acute and chronic colitis following *H.pylori* infection.

---

*Helicobacter pylori* (*H.pylori*) is the predominant bacterium colonized in the stomach, and associated with several diseases including gastritis, peptic ulcer, gastric cancer, and mucosa-associated lymphoid tissue (MALT) lymphomas(1). *H.pylori* infection can’t be cleared effectively even though it can elicit a series of immune responses. The immunological escape is considered to be mediated by the immunosuppressive regulatory cells. Foxp3 expression can be induced by *H.pylori* infection, and there is no or less Foxp3 mRNA expression in the stomach without *H.pylori* infection(2, 3). Gastritis caused by *H.pylori* is relieved by increased CD4+CD25+Foxp3+ marked regulatory T cells (Treg), which inhibit the Th1 and Th17 cells through the secretion of anti-inflammatory cytokines (IL-10、 TGF-β) and the manner of cell to cell contact (4, 5).

Our previous study showed that, besides the CD4^+^CD25^+^Foxp3^+^Treg cells, CD19+IL-10+ marked regulatory B cells (Breg) also expanded significantly after *H.pylori* infection. The regulatory immune responses induced by *H.pylori* infection were not limited to the stomach(6). In addition to Treg cells and Breg cells, regulatory dendritic cells are also associated with the immunological escape of *H.pylori* colonization(7, 8).

In recent years, more and more studies show that *H.pylori* infection is negatively associated with certain autoimmune diseases, such as inflammatory bowel disease (IBD), asthma, eczema, and so on(9–13).

IBD is a common kind of autoimmune diseases, consisting of ulcerative colitis (UC) and Crohn’s disease (CD), which are characterized as chronic and recurrent intestinal inflammation. The etiology and pathogenesis are undetermined, inappropriate activation of the mucosal immune system play an important role in the process of IBD(14, 15). Peter D.R.Higgins and colleagues(16) showed that *H.pylori* infection can reduce the Th17 immune response in cecum caused by Salmonella typhimurium infection, and increase the IL-10 expression in MLN. Jay Luther and colleagues(17) detected *H.pylori* DNA in intestine of *H.pylori* positive patients, which can inhibit the secretion of pro-inflammatory cytokine (IL-12、 IFN-I) from intestinal DC cells via TLR-9 pathway, alleviate the intestinal inflammatory responses induced by dextran sulphate sodium (DSS). It was also reported that *H.pylori* infection changed the balance of Th17/Treg cells and skewed to the Treg cell responses in intestinal DC cells, and then affected the intestinal mucosal immune responses(7, 18).

Numerous studies have shown that, experimental treatment of colitis induced by DSS was primarily performed by elevating the CD4^+^CD25^+^Foxp3^+^Treg cells, reducing the Th17 cells, increasing IL-10 expression and decreasing IL-1 β, IL-17 expression(15, 19). Ansary MM. et al found that apoptic cells could alleviated the chronic colitis induced by DSS through enhancing the function of Breg cells(20).

Induction of colitis by application of DSS in the drinking water is a widely used and well characterized model of colitis in mice(21). In our study, we use a rodent model, which was established acute and chronic colitis induced by DSS with or without *H.pylori* pre-infection, to investigate the effects of *H.pylori* infection on acute and chronic colitis, and the role of CD19^+^IL-10^+^Breg cells in the process.

## MATERIALS AND METHODS

### Animals

Female 6-8 week old C57BL/6 mice were purchased from Beijing Vitalriver Laboratory Animal Technology Company Limited, China. All mice were housed in specific pathogen-free rooms in microisolator cages in the Experimental Animal Center of Chinese Center for Disease Control and Prevention.

### *H.pylori* infection

All mice fasted overnight were intragastrically inoculated five times with 300 μl physiological saline containing 1×10^9^CFU/ml *H.pylori SS1* every other day. Age matched control mice were inoculated with physiological saline without *H.pylori SS1*.

Urease rapid test and Warthin-Starry were used to confirm the colonization of *H.pylori SS1*.

### DSS-induced colitis

Six weeks after *H.pylori* inoculated, 3% DSS (MP Biomedicals, CA, USA)was dissolved in sterile phosphate buffer and ad libitum for 7 days followed by substitution of phosphate buffer for another 7 days (one cycle) to induce acute colitis. Repeating the cycle three times to induce chronic colitis. All mice were assessed daily for body weight, diarrhea and bloody stool. The extent of the colitis was assessed by disease activity index (DAI). The DAI was calculated using the average of the total score: loss in body weigh (0, none; 1, 1-5%; 2, 5-10%; 3, 10-15%; 4, >15%); stool consistency (1, none; 2, loose stools; 4, diarrhea), and bloody stool (0, normal; 2, slight bleeding; 4, gross bleeding). There were five to six mice in each group.

### Flow cytometry

FITC-, APC-, PE– and PeyCP-cy5.5-conjugated anti-mouse CD4, CD25, Foxp3 (eBioscience, San Diego, CA, USA) and CD19, IL-10 (Biolegend, San Diego, CA) antibodies were used to stain the regulatory B cells and regulatory T cells. Briefly, after stimulating the lymphocytes with LPS and cell stimulation cocktail (plus protein transport inhibitors) (eBioscience) for 5 hours, CD19^+^IL-10^+^Breg cells were stained according to the manufacturer’s instruction of the Intracellular Fixation & Permeabilization Buffer set (Biolegend), including cell surface staining, Fixation & Permeabilization, and intracellular staining. CD4^+^CD25^+^Foxp3^+^Treg cells were stained according to the manufacturer’s instruction of Foxp3 staining buffer set (eBioscience). All samples were collected and analysied by BD FACS Calibur flow cytometer (BD Immunocytometry Systems, Franklin Lakes, NJ, USA). The date were analyzed by FlowJo 7.6 (FlowJo, Ashland, Oregon, USA).

### Histology and Immunohistochemistry

Sections of 4 μ m thick were cut from formalin-fixed and paraffin-embedded stomach and colon tissue and stained with hematoxylin and eosin (H&E) or Warthin-Starry. Histological scores were assessed according to L.A.Dieleman(14). Antigen retrieval was performed by heating the sections for 15min in citric acid solution (pH 6.0).Sections were incubated with rabbit anti-mouse Foxp3 antibody (1:200;Abcam,HongKong SAR,China) overnight at 4 °C after being blocked with sheep serum solution. Foxp3+ cells were quantified based on the mean number of stained cell in the lamina propria of two sections, including five different fields per section and excluding lymphoid follicles.

### Quantitative reverse-transcription polymerase chain reaction (qRT-PCR)

To detect the levels of the anti-inflammatory cytokines (IL-10、 TGFβ) and pro-inflammatory cytokines (IFNγ、 TNFα、 IL-17A、 IL-23) in the colonic mucosa, total RNA of colon was extracted using Trizol reagent (Ambion, Carlsbad, CA, USA) and reverse-transcribed to first-strand complementary DNA (cDNA) using the RevertAid First Strand cDNA Synthesis Kit (Thermo, Vilnius, Lithuania). RT-PCR was performed with a StepOne Plus Real-Time PCR System (Applied Biosystems, Waltham, MA, USA). Glyceraldehyde-3-Phosphate Dehydrogenase (GAPDH) was used as the internal control. Primers are listed in supplementary 1. All samples were analyzed in duplicate in a single 96-well reaction plate, and data were analyzed according to the 2^-ΔCt^ methods.

### Statistical analysis

All data were shown as mean ± SEM. Statistical analysis was performed with Statistical Package for Social Science version 17.0 software (SPSS Inc,Chicage.IL). Student’t t-test or one-way ANOVA was used to analyse values between different groups. P<0.05 was considered statistically significant. The graphs were ploted by GraphPad Prism 5 (GraphPad Software, La Jolla, CA, USA).

## Results

### *H.pylori* infection alleviate acute and chronic colitis induced by DSS

To investigate the effect of *H.pylori* infection on colitis induced by DSS, we treated wild type C57BL/6 mice, which infected with *H.pylori* SS1 six weeks ago, with 3% DSS for 7 days followed by PBS for 7 days as one cycle to induce acute colitis model, and repeated three cycles to induce chronic colitis model. The severity of colitis was evaluated by measuring DAI scores.

In acute colitis groups, body weight loss was first observed in DSS-treated mice on day 4. However, *H.pylori/* DSS co-treated mice began to display significant body weight loss on day 6. DSS-treated mice showed more weight loss between day 4 to day 8 compared with *H.pylori/* DSS co-treated mice (Figure 1A). DAI scores in DSS-treated mice were significantly increased on day 3 and higher than *H.pylori/* DSS co-treated mice between day 3 to 8(Figure 1B).The length of colon was mearsured when the mice sacrificed. It is also a parameter of the colitis severity. DSS-treated mice had shorter colon lengths than *H.pylori/* DSS co-treated mice(Figure 1C). To further assess colitis severity, the degree of colitis was evaluated histopathologically. DSS treatment induced eptithelial injury, crypt damage, lamina propria edema and thickened, and mononuclear cells infiltration transmural(Figure 1E). These changes were more severe in DSS-treated mice and the histology scores were In chronic colitis groups, the clinical performance of chronic colitis changed in keeping with the component in drinking water. The colon lengths of *H.pylori/* DSS co-treated mice shorten mild than DSS-treated mice, and colonic histology was significantly slight than DSS-treated mice (Figure S1). Thus, *H.pylori* infection can alleviate acute and chronic colitis induced by DSS.

**Figure 1.**
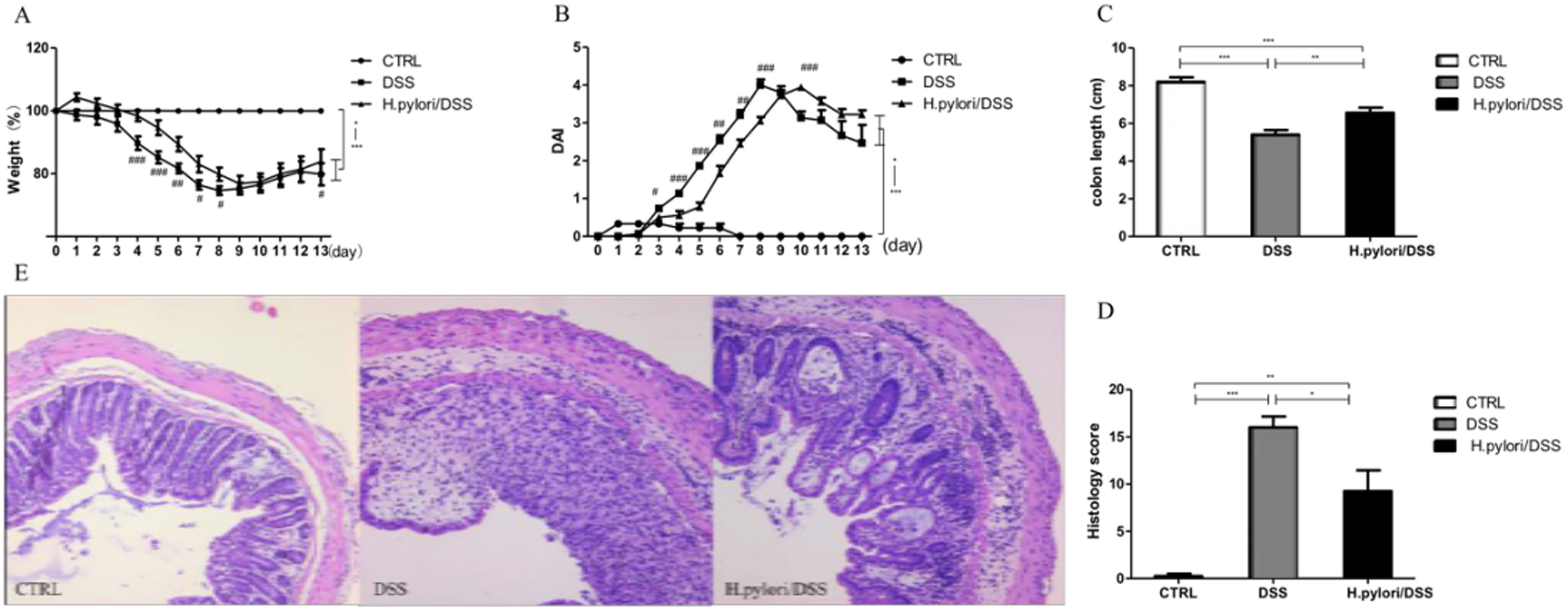
*H.pylori* infection alleviated acute colitis induced by DSS. A. Body weight of each group. B. Disease activation index (DAI) of each group. C. Colon length of each group. D. Histology score of each group. E. HE Staining of colon.(100X)(*,# P<0.05; **,## P<0.01; ***,###P<0.005. #,DSS vs *H.pylori*/DSS.)

### Gene expression of regulatory cytokine up-regulated while pro-inflammatory cytokine down-regulated in colon of *H.pylori/* DSS co-treated mice

Gene expression of cytokine reflect the immune and inflammatory status. We used quantitative real-time PCR to examine the expression of regulatory cytokine IL-10、 TGF-β and pro-inflammatory cytoine IFN-γ, TNF-α, IL-17A and IL-23 in colon.

In acute colitis groups, IL-10 and TGF-β mRNA expression levels up-regulated in DSS-treated mice and IL-10 mRNA was significantly lower than *H.pylori*/DSS co-treated mice (Figure 2A). Both IFN-γ and TNF-α mRNA expression levels up-regulated significantly in DSS-treated mice, while IFN-γ mRNA expression level was lower significantly in *H.pylori*/DSS co-treated mice (Figure 2A).

**Figure 2.**
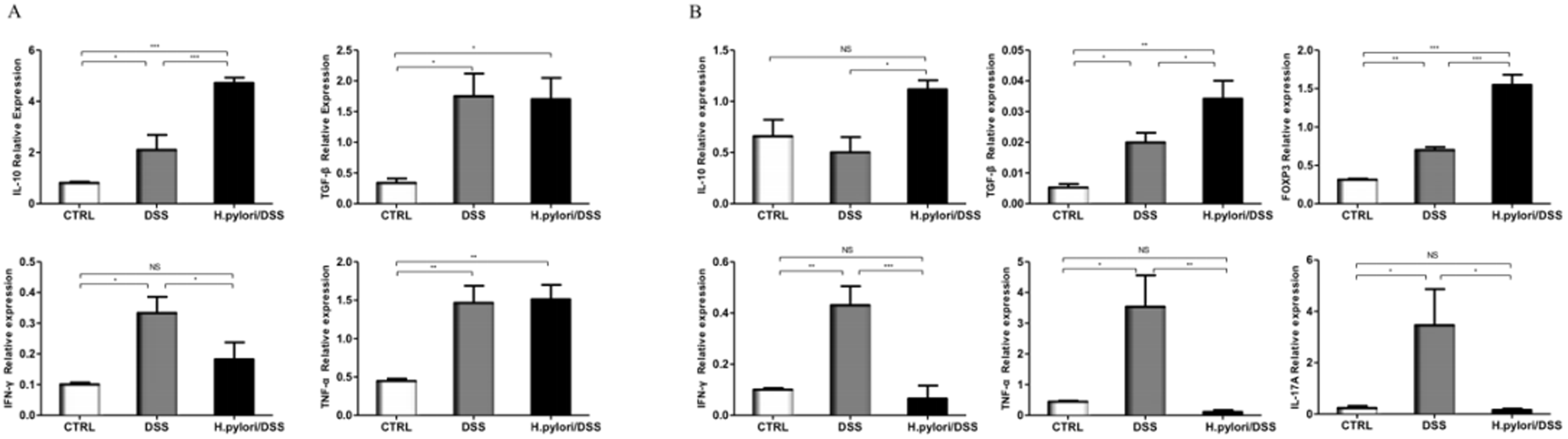
Gene expression of regulatory cytokine up-regulated with pro-inflammatory cytokine down-regulated in colon of *H.pylori*/DSS co treated mice. A. Cytokine expression in of acute colitis mice. B. Cytokine expression in colonn of chronic colitis mice. (*,P<0.05; **,P<0.01; ***,P<0.005; NS, no significant.)

In chronic colitis groups, IL-10, TGF-β and Foxp3 mRNA expression levels in colon were significantly higher in *H.pylori*/DSS co-treated mice comparing with DSS-treated mice (Figure 2B). The pro-inflammatory cytokine, including IFN-γ, TNF-α and IL-17A mRNA expression levels up-regulated significantly in DSS-treated mice, and significantly higher than *H.pylori*/DSS co-treated mice (Figure 2B). Thus, the regulatory and pro-inflammatory cytokine expression levels were consistent with the colonic histopathology.

### CD19^+^IL-10^+^Breg cells expanded notably in *H.pylori*/DSS co-treated mice

To determine whether CD19^+^IL-10^+^Breg cells play a role in the process of *H.pylori* infection affect DSS induced acute and chronic colitis, the percentages of CD19^+^IL-10^+^Breg cells were detected in peripheral blood, spleen, MLN and PP.

In acute colitis groups, CD19^+^IL-10^+^Breg cells in spleen, MLN, and PP, expanded in DSS-treated mice, and the percentages of CD19^+^IL-10^+^Breg cells in blood, spleen, MLN and PP were significantly higher in *H.pylori*/DSS co-treated mice(Figure 3B). The results in chronic colitis groups were similar to the acute colitis groups(Figure 3C). These data indicated that CD19^+^IL-10^+^Breg cells expanded in DSS induced acute and chronic colitis, and increased more notably in *H.pylori*/DSS co-treated mice.

**Figure 3.**
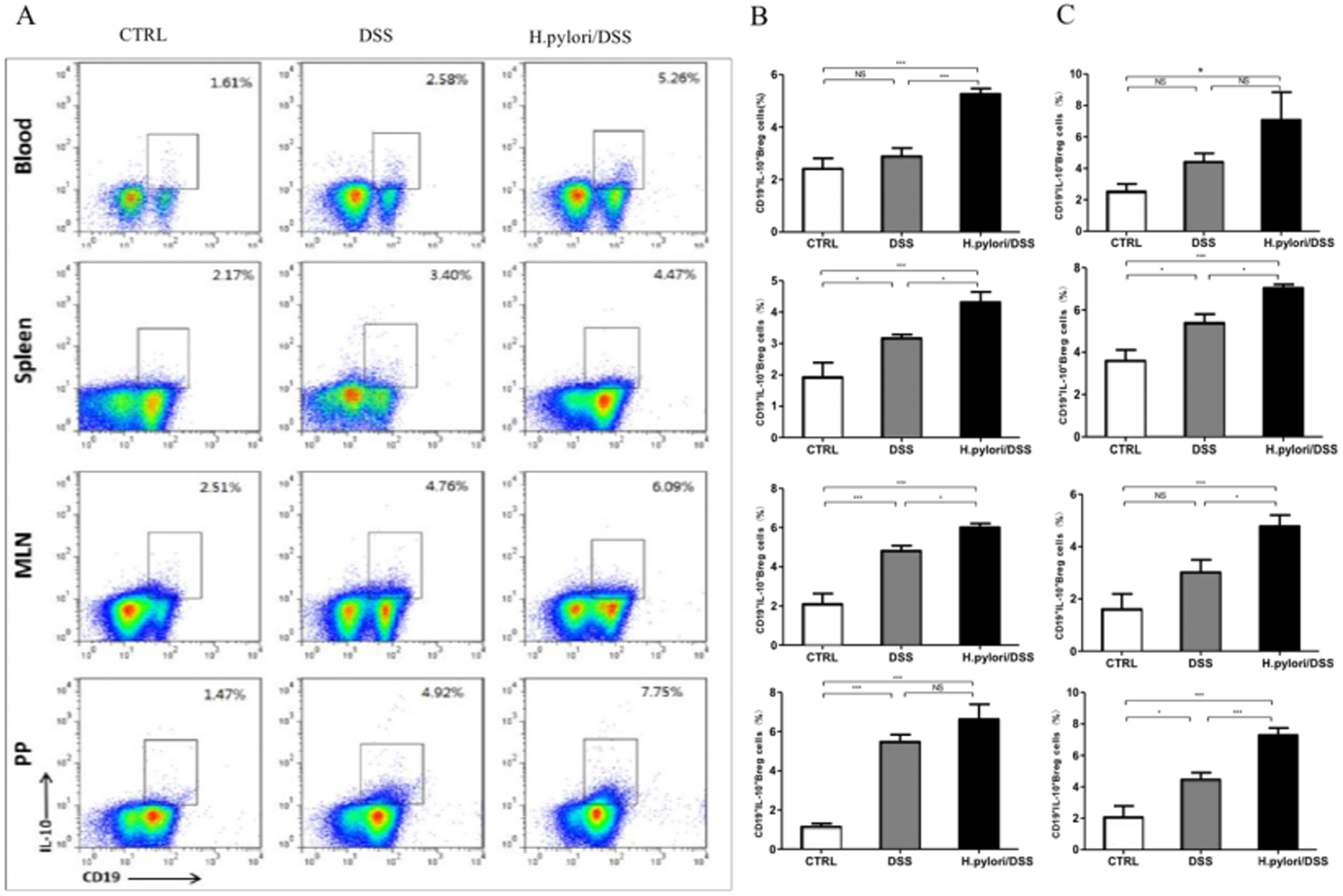
The percentages of CD19+IL-10+Breg cells increased notably in *H.pylori*/DSS co-treated mice in acute and chronic colitis mice. A. The percentages of CD19+IL-10+Breg cells in blood,spleen, MLN and PP were detected by flow cytometry. B. The percentages of CD19+IL-10+Breg cells in acute colitis mice. C. The percentages of CD19+IL-10+Breg cells in chronic colitis mice.(*,P<0.05; **,P<001; ***,P<0.005, NS, no significant.)

### CD4^+^CD25^+^Foxp3^+^Treg cells increased in acute colitis but decreased in chronic colits in *H.pylori*/ DSS co-treated mice

It is considered that, CD4^+^CD25^+^Foxp3^+^Treg cells are important in the development of IBD. In our study, we also detected the percentages of CD4^+^CD25^+^Foxp3^+^Treg cells in blood, spleen, MLN, and PP.

In acute colitis groups, the expansion of CD4^+^CD25^+^Foxp3^+^Treg cells were observed only in MLN in DSS-treated mice, and the percentages of CD4^+^CD25^+^Foxp3^+^Treg cells in spleen, MLN, and PP were significantly higher in *H.pylori*/DSS co-treated mice (Figure 4A). We also detected the Foxp3^+^Treg cells in colonic mucosa by immunohistochemistry. The results showed that Foxp3^+^Treg cells increased significantly in DSS-treated mice, and the extent of increase was more notable in *H.pylori*/DSS co-treated mice(Figure 4B).

**Figure 4.**
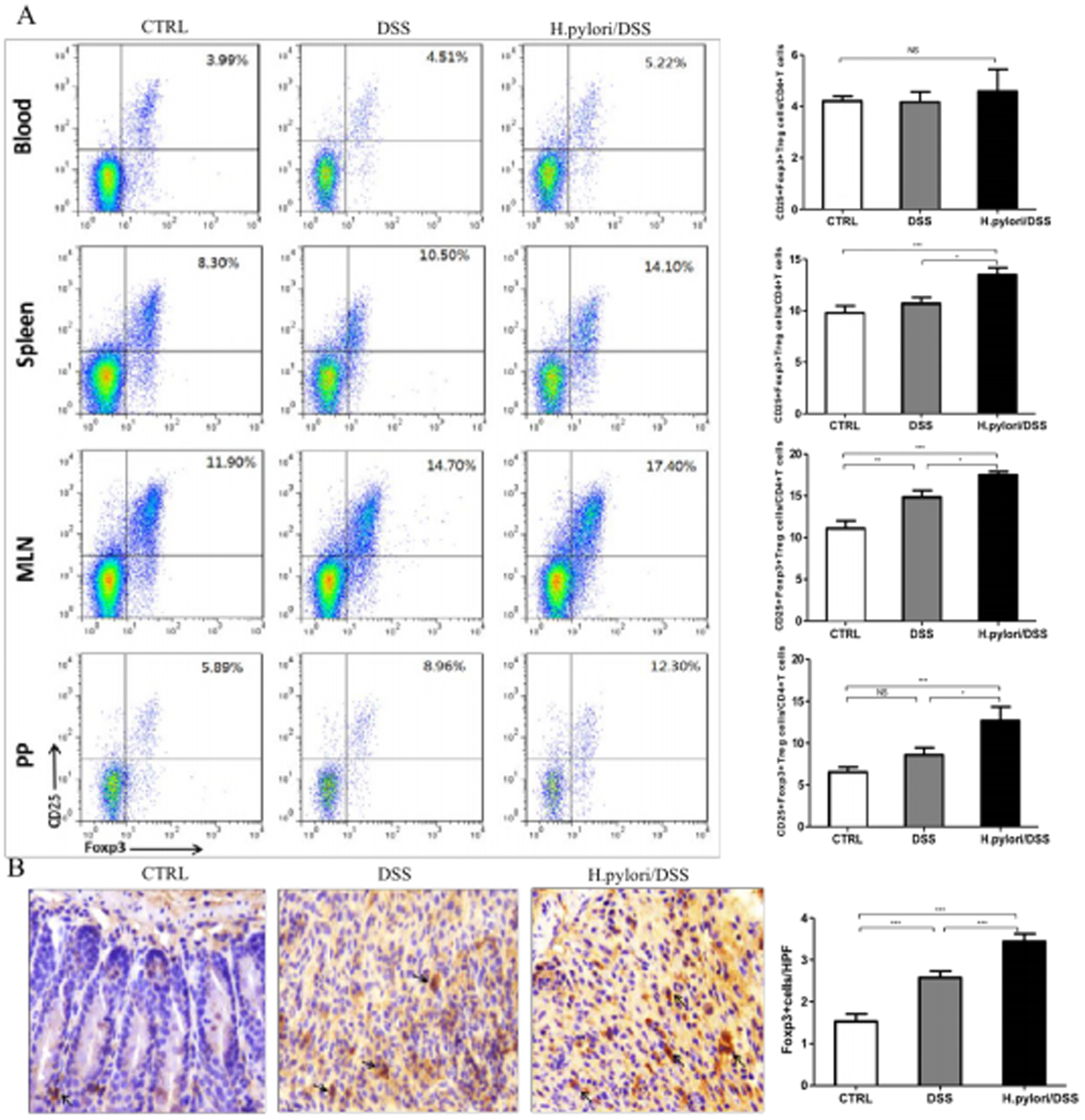
CD4+CD25+Foxp3+Treg cells increased in *H.pylori*/DSS co-treated acute colitis mice. A. CD4+CD25+Foxp3+Treg cells in blood,spleen,MLN and PP were detected by flow cytometry. B.Foxp3+Treg cells in colon were stained via immuno-histochemistry..(*,P<0.05; **,P<001; ***,P<0.005, NS, no significant.)

In chronic colitis groups, the changes of CD4^+^CD25^+^Foxp3^+^Treg cells were confusing, the percentages of CD4^+^CD25^+^Foxp3^+^Treg cells in blood, spleen, and PP were increased significantly, which was consistent with the acute colitis groups in DSS-treated mice. However, the percentages of CD4^+^CD25^+^Foxp3^+^Treg cells in tissues mentioned above were lower significantly in *H.pylori*/DSS co-treated mice(Figure 5A). The results of immunohistochemistry showed that there was no significant difference between the DSS-treated mice and *H.pylori*/DSS

**Figure 5.**
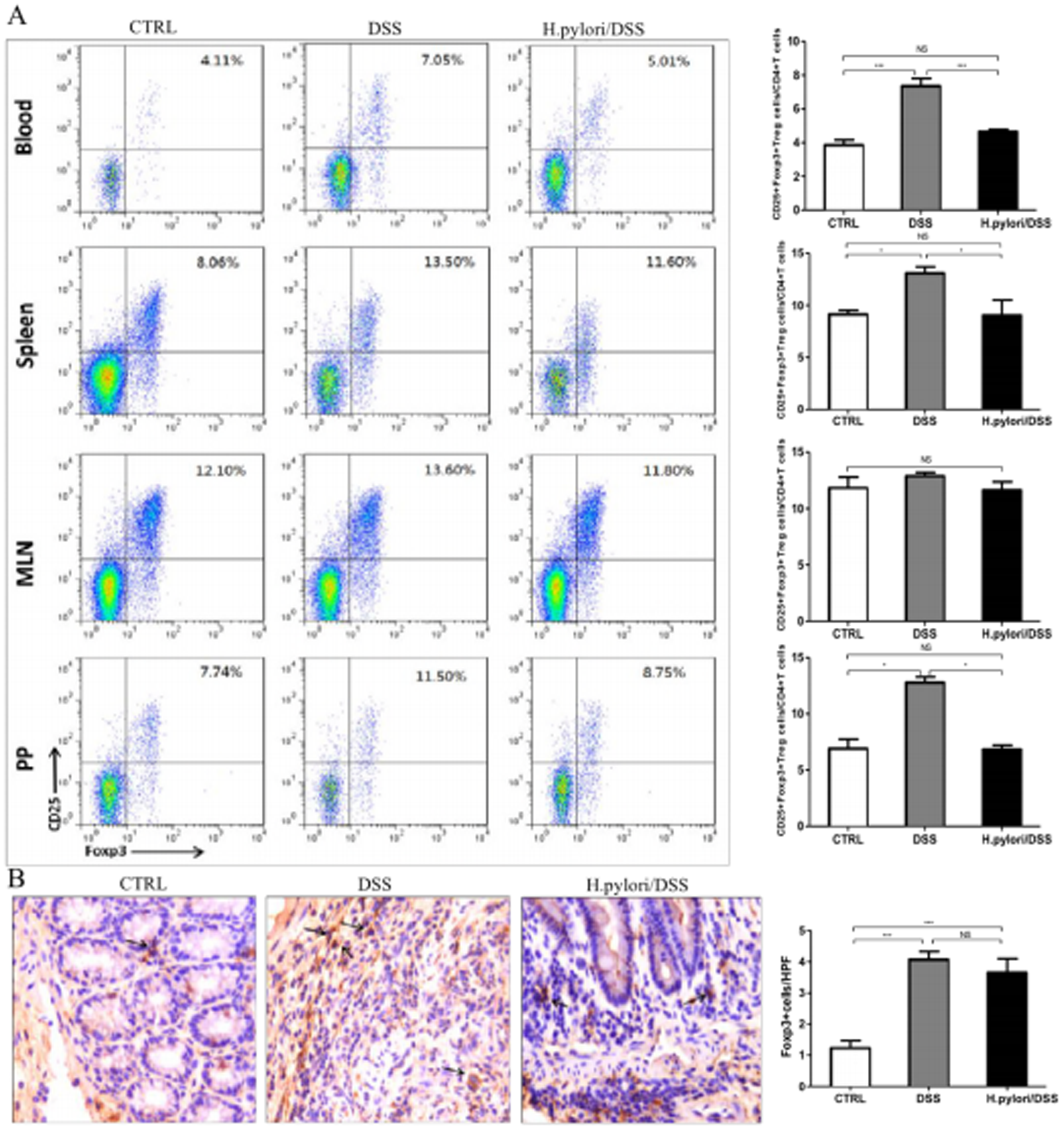
CD4+CD25+Foxp3+Treg cells degreased in *H.pylori*/DSS co-treated chronic colitis mice. A. CD4+CD25+Foxp3+Treg cells degreased significantly in blood,spleem and PP in *H.pylori*/DSS co-treated mice. B. There was no significant difference of Foxp3+Treg cells in colon between DSS-treated and *H.Tylori*/DSS co-treated mice. (*,P<0.05; **,P<001; ***,P<0.005, NS, no significant.)

## Disscussion

*H.pylori* infection is a world-wide popular question due to it’s contribution to the development of gastric cancer and peptic ulcer, and the infection will be lifelong without eradication. It was reported that the prevalence of IBD was low in *H.pylori* positive patients, and the objective of this study was to verify that *H.pylori* infection can ameliorate colitis, and investigate the function of Breg cells in the process. Given on the symptoms and histological characters and the cytokine expression in colon, the results support that *H.pylori* infection can alleviate acute and chronic colitis induced by DSS, which was consistent with epidemiologic studies(9, 22).

Breg cells were thought to be converted from different B cells stages in spleen firstly. Splenic B10 cells, as reported previously, expanded significantly during DSS induced acute colitis(23–26). It was revised that the conversion occurred in the inflammatory sites in recent years(27). Our previous study have showed that CD19^+^IL-10^+^Breg cells, ahead of CD4^+^CD25^+^Foxp3+Treg cells, expanded significantly in spleen, MLN, and gastrointestinal mucosa after *H.pylori* infection. Taking the previous studies into consideration, we detected the CD19^+^IL-10^+^Breg cells in different tissues including blood, spleen, MLN and PP in the rodent model. Our data showed that the percentages of CD19^+^IL-10^+^Breg cells both increased in acute and chronic colitis, and the extent of increase was more striking in acute and chronic colitis following *H.pylori* infection.

Breg cells can inhibit excessively activated immune response through the production of regulatory cytokines such as IL-10 and TGF-β. They can also co-operate with DC cells to promote the differentiation of Treg cells, resulting in the inhibition of effective T cells responses(28). So, we also detected the CD4^+^CD25^+^Foxp3+Treg cells in different tissues. In acute colitis groups, Treg cells increased mainly in the MLN in DSS-treated mice, and the percentages of Treg cells in *H.pylori*/DSS co-treated mice were significantly higher. This was also demonstrated that Breg cells expanded earlier than Treg cells. In chronic colitis groups, Tregs cells continued to expand in blood, spleen and PP. However, we surprisedly found that the percentages of Treg cells decreased significantly in *H.pylori*/DSS co-treated mice. We supposed that there were perhaps other regulatory cells such as immature DC cells or immunosuppressive macrophages to take part in the chronic colitis after *H.pylori* infection(29, 30).

There was an interesting discovery during the study that the numbers of PPs in the small intestine reduced and the volume diminished in DSS-treated mice. The numbers and volume of PPs recovered partly in *H.pylori*/DSS co-treated mice. Shigenori Nagai and colleagues thought that DC cells in PPs can recognize *H.pylori* antigen, initiate naive CD4^+^T cells differentiate into Th1 cells, which can migrate to the stomach and cause gastritis, *H.pylori* infection can activate systemic immune responses through PPs(31, 32). Spahn TW.et al reported that DSS can induce more severe colitis in PP-deficient mice compared with wild type mice, suggesting that PPs may protect the mice from the DSS(33). In summary, *H.pylori* may affect the acute and chronic colitis through regulating the function of PPs. It is to be confirmed and investigated through further experiments.

In conclusion, *H.pylori* infection can alleviate acute and chronic colits induced by DSS, and Breg cells play the more important role compared with Treg cells in the process. The inverse association between *H.pylori* infection and IBD provides a potential strategy and the Breg cells may be a promising target in IBD clinical treatment.

## Acknowledgements

This work was supported by the National Natural Science Foundation of China (81370515) and Beijing Natural Science Foundation (7162194). The authors thank He Lihua, National Institute for Communicable Disease Control and Prevention, Chinese Center for Disease Control and Prevention, for cultivating *H.pylori* SS1. No conflicts of interest exist.

